# Associations between thinner retinal neuronal layers and suboptimal brain structural integrity: Are the eyes a window to the brain?

**DOI:** 10.1101/2022.08.31.506114

**Authors:** Ashleigh Barrett-Young, Wickliffe C. Abraham, Carol Y. Cheung, Jesse Gale, Sean Hogan, David Ireland, Ross Keenan, Annchen R. Knodt, Tracy R. Melzer, Terrie E. Moffitt, Sandhya Ramrakha, Yih Chung Tham, Graham A. Wilson, Tien Yin Wong, Ahmad. R. Hariri, Richie Poulton

## Abstract

We investigated the extent to which measures of retinal neuronal thickness capture variability in the structural integrity of the brain in a large population-based cohort followed from birth to midlife. Using data from the Dunedin Multidisciplinary Health and Development Study (*n*=1037; analytic *n*=828, aged 45 years), we specifically tested for associations between optical coherence tomography-measured retinal neuronal layers and MRI-measured structural brain integrity. We found that Study members who had thinner retinal neuronal layers had thinner average cortex, smaller total cortical surface area, smaller subcortical grey matter volumes, larger volume of white matter hyperintensities as well as older looking brains. This suggests that retinal neuronal thickness reflects differences in midlife structural brain integrity consistent with accelerated cognitive decline and increased risk for later dementia, further supporting the proposition that the retina may be a biomarker of brain aging as early as midlife.

## 1. Introduction

The retina has potential as a biomarker of brain health and cognitive functioning as well as age-related neurodegenerative diseases, including Alzheimer’s disease (AD), because the retina shares many similarities with the brain and can be easily, repeatedly, and non-invasively imaged with high precision using technology which is already widely available (Alber et al., 2020; Cheung et al., 2021, 2017; London et al., 2013). Diffuse cortical atrophy occurs in normal ageing but is more pronounced in AD, beginning in the medial temporal lobes in preclinical AD and gradually progressing throughout the brain (Eckerström et al., 2018; Kulason et al., 2020; Pini et al., 2016). The neuropathological changes of AD begin more than a decade prior to the onset of clinical symptoms, (Sperling et al., 2014) and existing treatments for AD may be most effective in the earliest stages of the disease (Musiek and Morris, 2021). Thus, biomarkers that can identify people with preclinical AD or at high risk of developing AD, and that are able to be widely implemented for population screening, are imperative to initiating treatments at the optimal stage, preserving quality of life.

The retinal ganglion cells send their axons across the retinal surface to form the optic nerve, which projects posteriorly to targets in the brain (Chan et al., 2019; London et al., 2013). Normal ageing is associated with gradual thinning of the optic nerve; in AD a more profound loss of optic nerve axons can be found, in a diffuse and non-specific pattern, (Hinton et al., 1986) and this can be detected in early stages as thinning of the retinal nerve fibre layer (RNFL) and ganglion cell-inner plexiform layer (GC-IPL; Ge et al., 2021; Ikram et al., 2012; Mejia-Vergara et al., 2020). There is also specific amyloid detectable in retinal ganglion cell bodies and dendritic inputs, with some indication that the retina may be an early site for amyloid formation (Hadoux et al., 2019; Koronyo et al., 2012). There is evidence that subclinical retinal measures of optic nerve thinning occur in preclinical AD, and have been associated with indicators including amyloid burden, (Asanad et al., 2020; Ko et al., 2018; Santos et al., 2018), genetic risk (Santos et al., 2018), and cognitive performance (Barrett-Young et al., 2022; Ko et al., 2018).

A small number of studies have found associations between structural measurements of the brain with retinal measurements, although these studies, with the exception of the UK Biobank (Chua et al., 2021), have tended to involve older participants (≥ 65 years) or patients with diagnosed AD (Casaletto et al., 2017; den Haan et al., 2019; Donix et al., 2021; Jorge et al., 2020; Mejia-Vergara et al., 2021; Méndez-Gómez et al., 2018; Ong et al., 2015; Shi et al., 2019; Uchida et al., 2020; Ueda et al., 2022). The maximum clinical utility of retinal imaging in AD, however, is in the pre-diagnosis stages, before symptoms progress to the level where daily living is affected and irreversible neurological damage has occurred. Retinal imaging, being increasingly widely available even in primary care settings (e.g., optometry), as well as being repeatable and non-invasive, offers an attractive screening option for preclinical AD. Therefore, the question of interest is whether thinning of the retinal neuronal layers is associated with risk-related alterations in structural brain integrity in the decades before an AD diagnosis.

Recent studies have investigated these associations in cognitively unimpaired individuals, finding associations between retinal layers including the RNFL and GC-IPL and a number of structural brain measures, most commonly hippocampal volume (Chua et al., 2021; López-de-Eguileta et al., 2022; Shi et al., 2020), but also the lingual gyrus (Shi et al., 2020) and grey and white matter volumes (Chua et al., 2021). A recent study found correlations between RNFL, GC-IPL, and other retinal thickness measures with a large number of cortical thicknesses and volumes, in participants both with and without increased familial and genetic risk for AD (López-Cuenca et al., 2022). While most previous studies focus on elderly individuals, the current study contributes to the growing literature on this topic by examining a population-based sample of same-aged adults in midlife (aged 45 years at assessment), thus allowing us to examine the natural variations in the population without the effects of age. Our study design enables a unique opportunity to investigate early detectable changes in age-related neurological diseases in a sample currently free of AD diagnoses.

We hypothesised that thinner RNFL and GC-IPL measured by optical coherence tomography (OCT) would be associated with diminished structural brain integrity in midlife as reflected by thinner average cortex, smaller cortical surface area, smaller subcortical grey matter volumes and older looking brains as estimated using a machine learning-based estimate of chronological age based on MRI-derived measures (Liem et al., 2017). We also hypothesised that thinner RNFL and GC-IPL would be associated with a higher volume of white matter hyperintensities, a commonly used clinical marker of AD risk.

## 2. Method

### 2.1 Participants

Participants were members of the Dunedin Multidisciplinary Health and Development Study, a representative birth cohort (*n* = 1037; 91% of eligible births, 51.6% male) born between 1 April 1972 and 31 March 1973 in Aotearoa New Zealand. The cohort represents the full range of socioeconomic status in the general population of New Zealand and is predominantly New Zealand European (Pākehā; 93%). The study design and participant characteristics have been described extensively elsewhere (Poulton et al., 2015). Assessments were carried out at birth and ages 3, 5, 7, 9, 11, 13, 15, 18, 21, 26, 32, 38, and most recently at age 45 (2017-2019), when 94% of the 997 living Study members participated. The Dunedin Study was approved by the Health and Disability Ethics Committee, Ministry of Health, New Zealand. Study members gave informed consent before participating.

### 2.2 Optical coherence tomography (OCT)

OCT measurements were taken at age 45, between April 2017 and April 2019. OCT scans were performed in the morning by trained technicians using a spectral domain OCT machine (Cirrus HD-OCT, model 5000; Carl Zeiss Meditec). Mean peripapillary retinal nerve fibre layer (RNFL) and mean macular ganglion cell-inner plexiform layer (GC-IPL) thickness were calculated. RNFL thicknesses were generated on a 3.5mm circle from the optic disc cube. GC-IPL was generated from macular cube scan. Trained graders checked all scans for quality. Scans were removed from the final dataset due to insufficient OCT image quality (e.g. signal strength below 6, scan not correctly positioned, or image artefacts). Seven Study members were removed due to diseases affecting the retina (multiple sclerosis, retinitis pigmentosa, brain tumours, diabetic laser pan-retinal photocoagulation, and an anomalous optic nerve head). Another seven Study members were assessed by two ophthalmologists as having glaucoma (Singh et al., 2022); glaucomatous eyes were removed from the dataset and non-glaucomatous eyes were retained. When data from one eye were available, that eye was used; when both eyes were available, an average of the measurements from both eyes was used. Axial length was used as a covariate as it influences RNFL and GC-IPL thickness measurements (Röck et al., 2014). Axial length was measured for right and left eyes using Zeiss IOL Master (Germany). The machine was calibrated weekly, with room lighting set at 520 lux and no pupil dilation. Mean axial length was calculated from both eyes.

### 2.3 Magnetic resonance imaging (MRI)

Study members were scanned using a MAGNETOM Skyra 3T scanner (Siemens Healthcare, Erlangen, Germany) equipped with a 64-channel head and neck coil (due to head size constraints, seven participants were scanned with a 20-channel head/neck coil) at the Pacific Radiology Group imaging centre in Dunedin, New Zealand, between August 2016 and April 2019. High resolution T1-weighted images, three-dimensional fluid-attenuated inversion recovery (FLAIR) images, and a gradient echo field map were obtained. Structural MRI data were analysed using the Human Connectome Project (HCP) minimal preprocessing pipeline (Glasser et al., 2013). Outputs of the preprocessing pipeline were visually checked for accurate surface generation by examining each participant’s myelin map, pial surface, and white matter boundaries. Study personnel who processed the MRI images were masked to participants’ retinal measurements. Of the participants with available structural MRI data, four were excluded due to major incidental findings or previous injuries (e.g. tumour or extensive damage to brain or skull), nine due to missing FLAIR or field map scans, and one due to poor surface mapping. Additionally, white matter hyperintensities measurements were removed from the dataset for three SMs due to multiple sclerosis and 8 Study members due to inaccurate labelling or low-quality data.

The brain Age Gap Estimate (brainAGE) is a score that represents the difference, or gap, between a person’s chronological age and their estimated age based on multiple measures of brain structure including cortical thickness, surface area, and volume of subcortical grey matter, white matter, and cerebrospinal fluid (Liem et al., 2017). The formation of this measure has been described previously (Elliott et al., 2019). White matter hyperintensities (WMH) were identified and extracted from T1-weighted and FLAIR images processed using UBO Detector, a cluster-based, fully-automated pipeline (Jiang et al., 2018). Grey matter volumes were extracted for 10 subcortical structures using the FreeSurfer aseg parcellation (https://surfer.nmr.mgh.harvard.edu/). Mean total cortical surface area and average cortical thickness were estimated over 360 cortical areas in the HCP-Multi Modal Parcellation version 1.0 (Glasser et al., 2016).

#### 2.3.1 Image acquisition parameters

High resolution T1-weighted images were obtained using an MP-RAGE sequence with the following parameters: TR = 2400 ms; TE = 1.98 ms; 208 sagittal slices; flip angle, 9°; FOV, 224 mm; matrix = 256×256; slice thickness = 0.9 mm with no gap (voxel size 0.9×0.875×0.875 mm); and total scan time = 6 min and 52 s. 3D fluid-attenuated inversion recovery (FLAIR) images were obtained with the following parameters: TR = 8000 ms; TE = 399 ms; 160 sagittal slices; FOV = 240 mm; matrix = 232×256; slice thickness = 1.2 mm (voxel size 0.9×0.9×1.2 mm); and total scan time = 5 min and 38 s. Additionally, a gradient echo field map was acquired with the following parameters: TR = 712 ms; TE = 4.92 and 7.38 ms; 72 axial slices; FOV = 200 mm; matrix = 100×100; slice thickness = 2.0 mm (voxel size 2 mm isotropic); and total scan time = 2 min and 25 s.

#### 2.3.2 Image processing

Structural MRI data were analyzed using the Human Connectome Project (HCP) minimal preprocessing pipeline as extensively detailed elsewhere (Glasser et al., 2013). Briefly, T1-weighted and FLAIR images were processed through the PreFreeSurfer, FreeSurfer, and PostFreeSurfer pipelines. T1-weighted and FLAIR images were corrected for readout distortion using the gradient echo field map, coregistered, brain-extracted, and aligned together in the native T1 space using boundary-based registration (Greve and Fischl, 2009). Images were then processed with a custom FreeSurfer recon-all pipeline that is optimized for structural MRI with higher resolution than 1 mm isotropic. Finally, recon-all output were converted into CIFTI format and registered to common 32k_FS_LR mesh using MSM-sulc (Robinson et al., 2014).

For each subject the mean cortical thickness and surface area were then extracted from each of the 360 cortical areas in the HCP-MPP1.0 parcellation (Glasser et al., 2016). Subcortical volumes were extracted separately using the automatic segmentation (“aseg”) step of FreeSurfer version 6.0. FreeSurfer version 6.0 was used because the HCP FreeSurfer pipeline was optimized for the cortical surface, resulting in lower-quality segmentation of subcortical volumes in our dataset. Outputs of the minimal preprocessing pipeline were visually checked for accurate surface generation by examining each subject’s myelin map, pial surface, and white matter boundaries. Accuracy of subcortical segmentation was confirmed by visual inspection of the “aseg” labels overlaid on the volumes. Of the 875 study members for whom data were available, 4 were excluded due to major incidental findings or previous injuries (e.g., large tumours or extensive damage to the brain/skull), 9 due to missing FLAIR or field map scans, and 1 due to poor surface mapping, yielding 861 datasets for analyses.

To identify and extract the total volume of white matter hyperintensities (WMH), T1-weighted and FLAIR images for each participant were processed with UBO Detector, a cluster-based, fully-automated pipeline with high reliability in our data (test-retest ICC = 0.87, 95% CI = [.73, .95]) and out of sample performance (Jiang et al., 2018). The resulting WMH probability maps were thresholded at 0.7, which is the suggested standard. WMH volume is measured in Montreal Neurological Institute (MNI) space, removing the influence of differences in brain volume and intracranial volume on WMH volume. Because of the potential for bias and false positives due to the thresholds and masks applied in UBO, the resulting WMH maps for each participant were manually checked by two independent raters to ensure that false detections did not substantially contribute to estimates of WMH volume. Visual inspections were done blind to the participants’ cognitive status. Due to the tendency of automated algorithms to mislabel regions surrounding the septum as white matter hyperintensities, these regions were manually masked out, to further ensure the most accurate grading possible. Of the 875 Study members for whom WMH data were available, 4 were excluded due to major incidental findings or previous injuries (e.g., large tumors or extensive damage to the brain/skull), 8 due to missing FLAIR scans, 3 due to diagnosis with multiple sclerosis, and 8 due to inaccurate white matter labelling or low-quality MRI data, yielding 852 datasets for analyses.

### 2.4 Data analysis

Analyses were conducted in Stata/SE 17.0 between February and July 2022. First, linear regression models were constructed where each retinal variable was entered as a predictor of brainAGE (ICC for test-retest reliability = .81); next, mean RNFL and mean GC-IPL were used to predict global structural measures (total cortical surface area, average cortical thickness, and white matter hyperintensities volume; ICCs = .996, .94, and .87 respectively). White matter hyperintensities volume was log transformed as the data were non-normal; all other variables met criteria for parametric testing without adjustment. Next, we used mean RNFL and mean GC-IPL to predict the grey matter volume (GMV) of 10 subcortical structures (mean ICC = .956). Finally, to explore the distribution patterns of associations, parcel-wise analyses were conducted when statistically significant associations were observed with global cortical surface area and average cortical thickness. In these post-hoc analyses, we ran linear regressions using mean RNFL and mean GC-IPL to predict the surface area and cortical thickness each of the 360 regions comprising the parcellation scheme described above (Glasser et al., 2016; mean ICCs = .846 and .942 for parcel-wise cortical thickness and surface area, respectively). As mean RNFL was not significantly associated with average cortical thickness, parcel-wise analysis with mean RNFL was only conducted with surface area. We corrected for multiple comparisons across the subcortical and parcel-wise analyses performed using a false discovery rate (FDR) procedure (Benjamini and Hochberg, 1995); for analyses with global variables we used an alpha level of .05. All tests were two-sided. Sex and axial length were included as covariates in all analyses. Analyses with regional subcortical structures were repeated controlling for total brain volume, which tests relative size of a region rather than absolute size (Hariri, 2020). The premise and analysis plan for this study was preregistered in December 2021 (https://dunedinstudy.otago.ac.nz/files/1639953449_Barrett-Young_CP_OCT%20and%20MRI_revised_final.pdf). Analyses were checked for reproducibility by an independent data analyst, who recreated the code by working from the manuscript and applying it to a fresh dataset.

## 3 Results

Data collection of retinal and brain structural data was at 45 years of age, completed between August 2016 and April 2019. The final dataset included those Study members with both RNFL and MRI data available (*n* = 828, female *n* = 413 [49.9%], male *n* = 415 [50.1%]) for analyses using RNFL variables, and those Study members with both GC-IPL and MRI data available (*n* = 825, female *n* = 413 [50.1%], male *n* = 412 [49.9%]) for analyses using GC-IPL variables. See Figure 1 for diagram of retinal measures and Table 1 for descriptive statistics of retinal and MRI measures.

**Table 1.** Descriptive statistics.

|  | *n* | *M* | *SD* |
| --- | --- | --- | --- |
| Mean RNFL thickness (μm) | 863 | 92.83 | 9.31 |
| Mean GC+IPL Thickness (μm) | 859 | 82.87 | 5.86 |
| brainAGE (z score) | 869 | 0.00 | 8.01 |
| Cortical surface area (mm<sup>2</sup>)<sup>a</sup> | 861 | 185472.12 | 16347.12 |
| Cortical thickness (mm)<sup>a</sup> | 861 | 2.56 | .09 |
| White matter hyperintensities<sup>a,b</sup> | 852 | 6.52 | .80 |
| Accumbens volume (mm<sup>3</sup>) | 861 | 488.25 | 77.81 |
| Amygdala volume (mm<sup>3</sup>) | 861 | 1739.80 | 204.71 |
| Brain stem volume (mm<sup>3</sup>) | 861 | 21775.42 | 2564.87 |
| Caudate volume (mm<sup>3</sup>) | 861 | 3313.65 | 421.36 |
| Cerebellum cortex volume, (mm<sup>3</sup>) | 861 | 60482.18 | 6250.09 |
| Hippocampus volume (mm<sup>3</sup>) | 861 | 4322.67 | 423.3 |
| Pallidum volume (mm<sup>3</sup>) | 861 | 1976.91 | 215.94 |
| Putamen volume (mm<sup>3</sup>) | 861 | 4699.70 | 536.45 |
| Thalamus volume (mm<sup>3</sup>) | 861 | 7785.12 | 797.95 |
| Ventral diencephalon volume (mm<sup>3</sup>) | 861 | 4141.99 | 409.63 |

<sup>a</sup>Whole brain; <sup>b</sup>log transformed.

**Figure 1.**
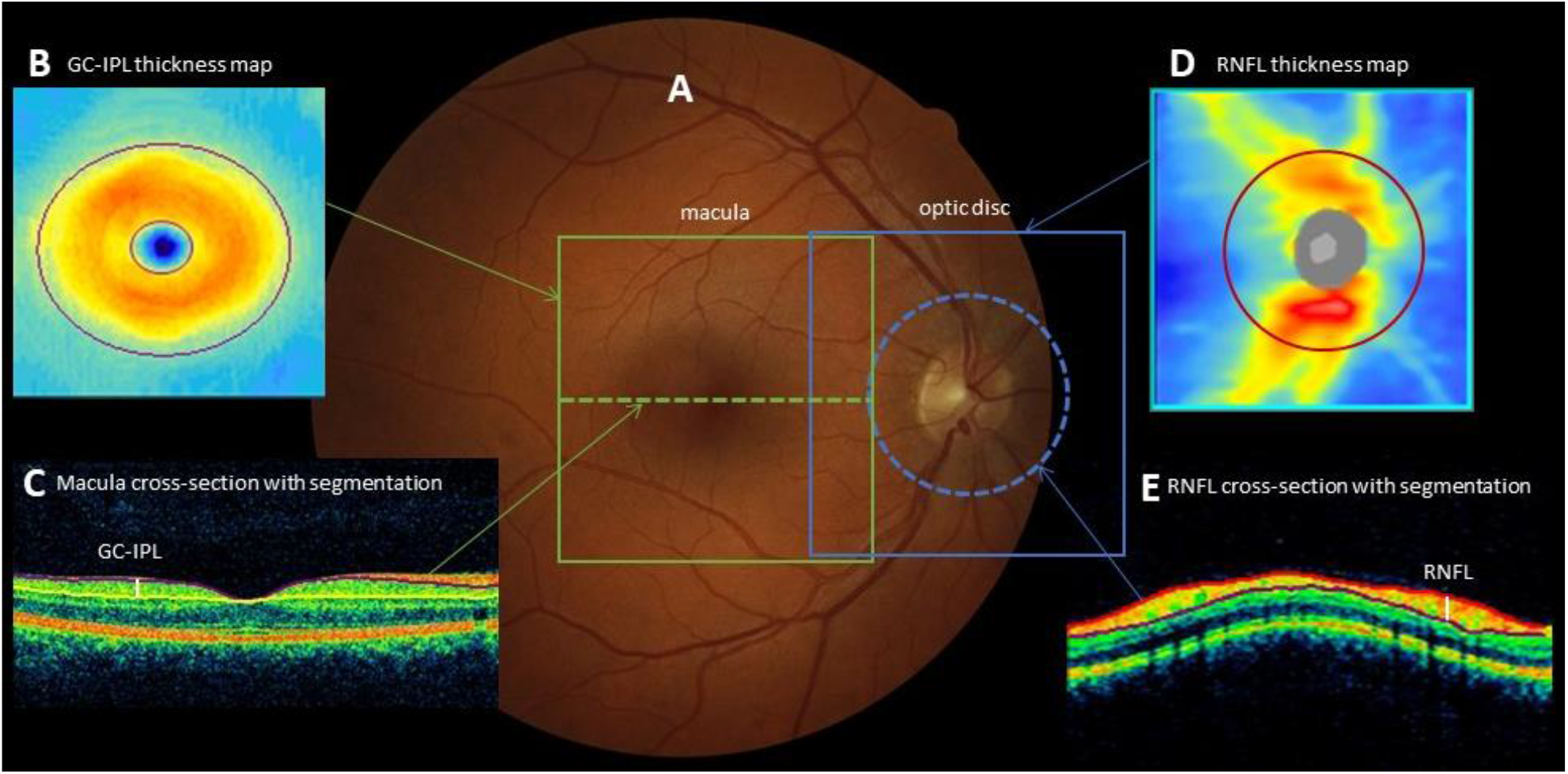
Retinal measurements of optic nerve structure. (A) Fundus photograph of a normal eye showing the areas scanned. (B) Ganglion cell-inner plexiform layer (GC-IPL) map showing a normal pattern. The GC-IPL thickness used for analysis was the average across the donut shaped area between the two ovals. (C) OCT cross-section through the fovea (indicated on main photo by dotted line), with automated segmentation lines showing the GC-IPL layer thickness measurement (indicated by the white line). (D) The retinal nerve fibre layer (RNFL) map shows thickest part of the nerve fibre layer close to the optic disc and at the superior and inferior poles. The thickness is calculated on a circle of 3.5mm diameter (indicated by red line, and on main photo by dotted line). (E) The retinal cross section on the 3.5mm circle, showing automated segmentation and the RNFL thickness measurement (indicated by white line). The RNFL thickness used for analysis was the average thickness around the 3.5mm circle.

### 3.1 Retinal predictors of brain age and cortical brain structure

Study members with thinner mean RNFL (β = -.119, *p* < .001) and thinner mean GC-IPL (β = -.111, *p* = .001) had a larger brain age gap estimate (brainAGE; Liem et al., 2017) after adjustment for sex and axial length, indicating an older looking brain (Figure 2). Study members with thinner mean RNFL (β = .140, *p* < .001) and thinner mean GC-IPL (β = .095, *p* < .001) had smaller total cortical surface area. Study members with thinner mean GC-IPL (β = .072, *p* = .043) but not mean RNFL (β = .036, *p* = .328) had thinner average cortex. Study members with thinner mean GC-IPL (β = -.071, *p* =.045) but not mean RNFL (β = -.068, *p* = .061) had a larger volume of white matter hyperintensities.

**Figure 2.**
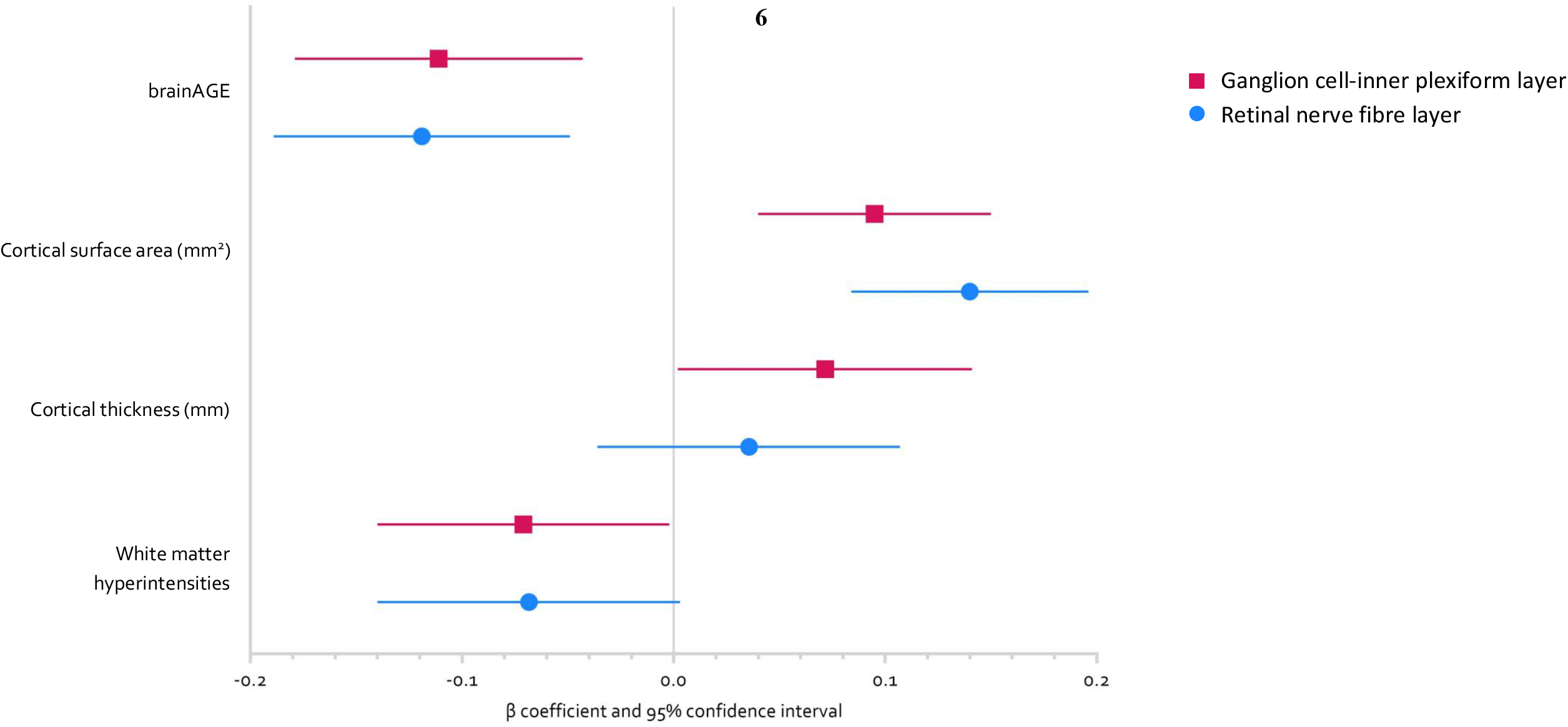
Forest plot showing standardised effect sizes (beta coefficients with 95% confidence intervals) for the associations between retinal nerve fibre layer (RNFL; blue circles) and ganglion cell-inner plexiform layer (GCL; pink squares) and whole brain variables. All models were adjusted for sex and axial length. Number of observations for each analysis differs due to differing quality control criteria (brainAGE and RNFL *n* = 828; cortical SA/CT and RNFL *n* = 821; WMH and RNFL *n* = 814; brainAGE and GC-IPL *n* = 825; cortical SA/CT and GC-IPL *n* = 818; WMH and GC-IPL *n* = 812. SA = surface area; CT = cortical thickness; WMH = white matter hyperintensities (log transformed)).

### 3.2 Retinal predictors of subcortical grey matter volume (GMV)

Study members with thinner mean RNFL and mean GC-IPL had smaller GMV of all ten subcortical structures after false discovery rate correction, indicating that retinal thinning corresponds to non-localised, rather than specific regional, subcortical GMV loss (Figure 3). Adding total brain volume as a covariate attenuated the associations between some subcortical GMVs and RNFL and GC-IPL, but associations remained significant between both retinal variables and GMV of brain stem, cerebellum, hippocampus, pallidum, and ventral DC; as well as between RNFL and GMV of the putamen and thalamus (Figure 3).

**Figure 3.**
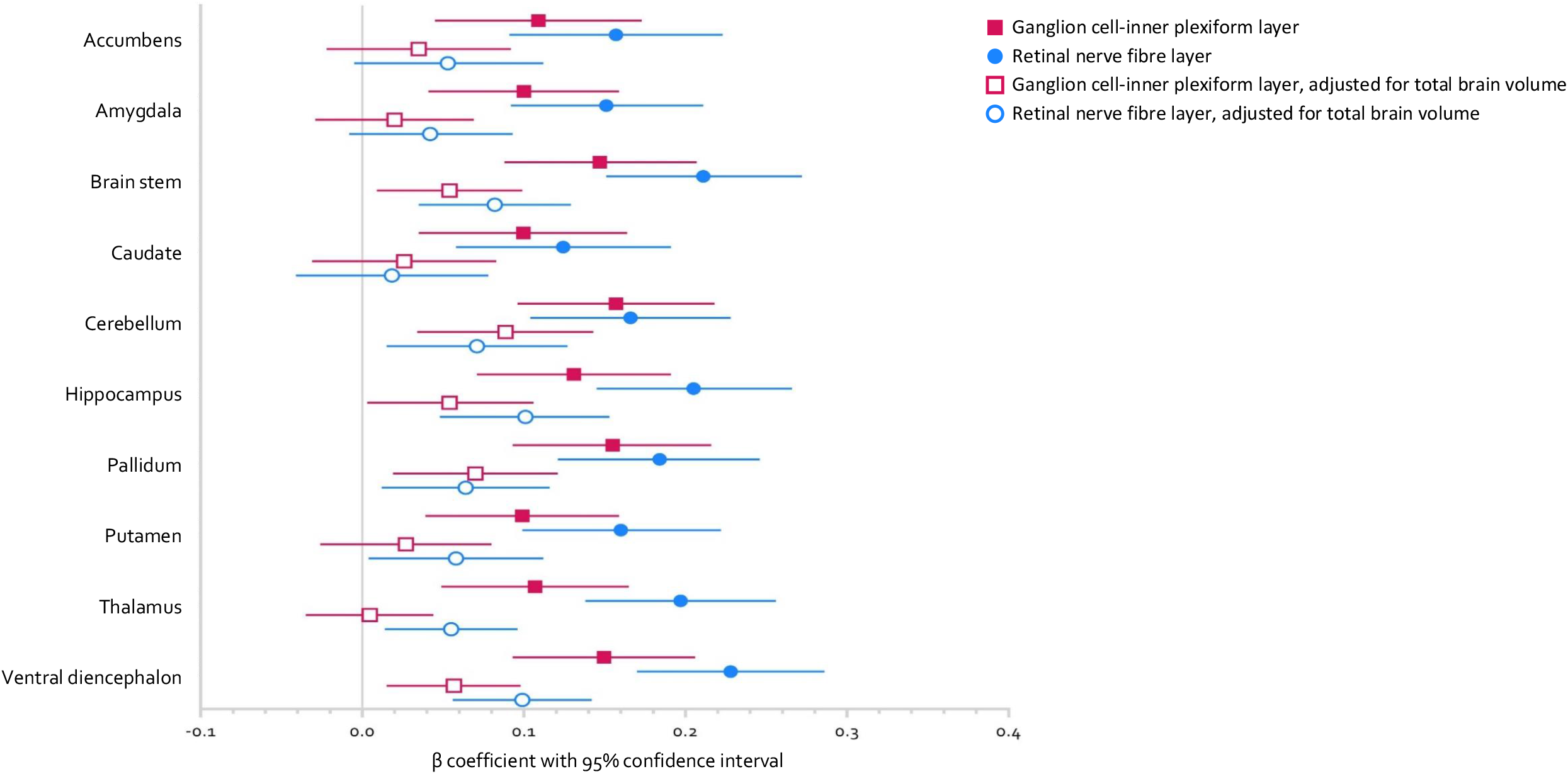
Forest plot showing standardised effect sizes (beta coefficients with 95% confidence intervals) for the associations between retinal nerve fibre layer (RNFL; blue circles) and ganglion cell-inner plexiform layer (GCL; pink squares) and grey matter volume of ten subcortical regions. Closed shapes indicate model was adjusted for sex and axial length; open shapes indicate model was adjusted for total brain volume as well as sex and axial length. A false discovery rate procedure was used to correct for multiple comparisons. Number of observations for analyses with RNFL *n* = 821; GCL *n* = 818. Ventral DC = ventral diencephalon.

### 3.3 Exploratory parcel-wise analysis

Based on significant associations observed between RNFL and GC-IPL and total cortical surface area, and GC-IPL and average cortical thickness, exploratory parcel-wise analyses were conducted to determine whether the associations were largely distributed or localised (Figure 4). Study members with thinner RNFL had lower surface area in 274 of 360 parcels after correction for multiple comparisons, which were widely distributed across the cortex. Study members with thinner GC-IPL had lower surface area in 112 parcels, again widely distributed across the cortex. Study members with thinner GC-IPL had thinner cortex in 5 parcels, which tended to be localised to the occipital and temporal lobes but were not widely distributed across these lobes.

**Figure 4.**
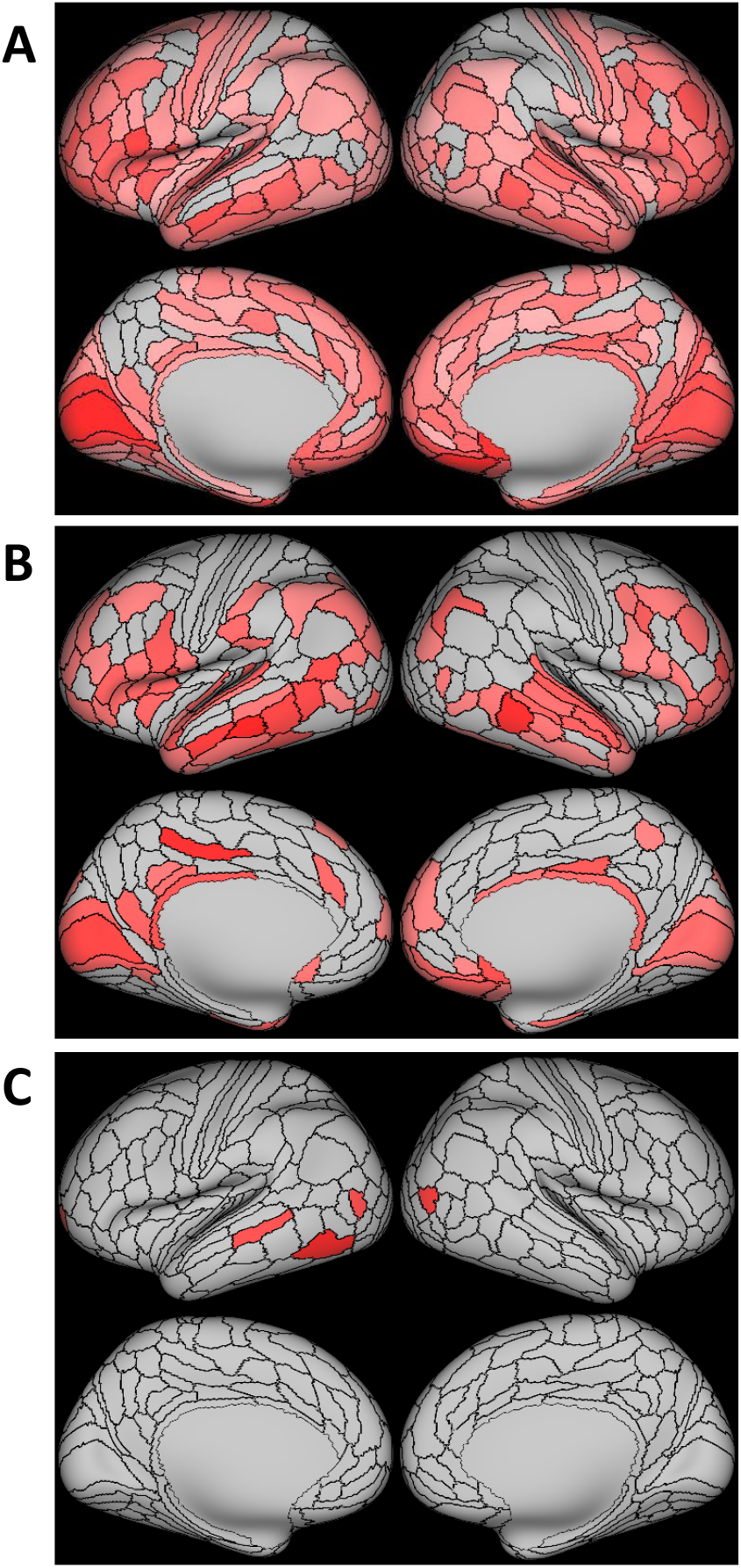
Associations between retinal thickness and parcel-wise measures of cortical surface area and thickness. (A) Retinal nerve fibre layer thickness and cortical surface area (*n* = 821); (B) Ganglion cell-inner plexiform layer thickness and cortical surface area (*n* = 818); (C) Ganglion cell-inner plexiform layer thickness and cortical thickness (*n* = 818). All models were adjusted for sex and axial length. Coloured parcels represent standardised coefficients with *p* < .05 after false discovery rate correction.

## 4 Discussion

We found that thinner retinal neuronal layers measured by OCT were associated with suboptimal MRI-measured brain structure and integrity, in a sample of largely healthy community-based middle aged people. Specifically, thinner RNFL and GC-IPL were associated with diminished structural brain integrity in midlife, as well as lower total cortical surface area and lower subcortical GMV and older looking brains. Study members with thinner GC-IPL, but not RNFL, also had thinner average cortex and larger volume of white matter hyperintensities. Exploratory analyses of parcel-wise surface area and thickness revealed that associations with RNFL and GC-IPL were widely distributed across the cortex, rather than regionally-specific. These findings provide supporting evidence that the retinal neuronal layers reflect alterations found in midlife structural brain integrity associated with later risk for AD, supporting the proposition that the retina may be a useful biomarker for brain health as early as midlife, and that non-invasive retinal imaging using OCT may potentially be useful to detect early preclinical disease.

An increasing number of studies have investigated whether retinal neuronal measurements are associated with brain measurements at various stages of AD. The findings are inconsistent, with some studies finding an association between retinal and brain measurements (Jorge et al., 2020; López-de-Eguileta et al., 2022; Mauschitz et al., 2022; Mejia-Vergara et al., 2021; Méndez-Gómez et al., 2018; Mutlu et al., 2018, 2017; Shi et al., 2020, 2019; Uchida et al., 2020), some finding no association (Casaletto et al., 2017; den Haan et al., 2019), and others with inconclusive results (Donix et al., 2021). However, methodological differences may explain some of this inconsistency—systematic differences between OCT or MRI devices across manufacturers and models, choice of parameters and settings, and the wide range of measurements which can be acquired from each of these technologies make comparison across studies difficult. In particular, some studies have focused on the hippocampus as an a priori region of interest, while others have taken a more global approach by assessing whole-brain, regional, or voxel-based measures.

One of the largest studies to investigate this question to date, the Singapore Epidemiology of Eye Diseases Study, found that thinner GC-IPL was associated with reduced total brain volume and GMV in two out of five lobes, but thinner RNFL was associated with GMV in the temporal lobe only (Ong et al., 2015). This aligns with our findings, where GC-IPL was associated with a larger number of MRI measurements than RNFL, although different regions were evaluated in each study. Another large study found that RNFL and GC-IPL thickness were associated with grey and white matter measurements in the visual pathway, but not globally (Mutlu et al., 2018). This is somewhat contradictory to our findings, which suggest a more global association pattern, but we note that when cortical thickness was examined at the parcel-level, the largest associations with GC-IPL thickness tended to be in the occipital and temporal lobe, including the visual cortex. A recent study from the UK Biobank suggests that retinal measures are associated with smaller cortical and hippocampal volume in a large middle-aged volunteer sample (Chua et al., 2021). However, this sample is biased in favour of higher education, socioeconomic status, and lower levels of adversity; our findings thus add to those of the UK Biobank by examining a more widely-representative, population-based sample (Brayne and Moffitt, 2022).

Retinal imaging technologies have a huge potential as a biomarker and predictive tool in preclinical AD, particularly with the development of artificial intelligence approaches (Ng et al., 2021). There has been rapid development of hardware (e.g., cheaper, higher resolution, and faster scanning speed OCT devices) and software (e.g. artificial intelligence algorithms), for retinal imaging. Recent advances have included reliable diagnosis of optic disease and risk stratification of morbidity and mortality using artificial intelligence and deep learning (Milea et al., 2020; Nusinovici et al., 2022). However, retinal thinning is not specific for AD and may occur in ageing and in other neurodegenerative diseases. This study provides evidence for links between at-risk patterns of structural brain integrity and retinal thinning in midlife, decades before any AD diagnosis. The retina is likely to contain more numerous and specific biomarkers of ageing and AD, such as vascular abnormalities, angiography, thinning of other retinal layers that can be captured with emerging technologies such as hyperspectral or amyloid imaging of the retina. Combining these retinal biomarkers and other risk factors to predict risk could leverage an artificial intelligence or machine learning approach, to both integrate and differentiate these risk factors for accelerated ageing and AD (Ng et al., 2021).

It is notable that between-subjects differences in both brain and retinal health were evident in this sample of middle-aged participants who were all the same chronological age. This suggests that anatomical changes typical of age-related diseases are detectable in midlife, decades before any expected symptom onset or diagnosis. Structural brain alterations, including white matter hyperintensities and older looking brains, have been associated with cognitive decline from childhood to age 45 in the current sample (d’Arbeloff et al., 2019; Elliott et al., 2019), as has retinal thinning (Barrett-Young et al., 2022), suggesting that neuropathological processes are associated with subclinical cognitive decline decades before typical AD diagnosis. Accelerated thinning of the optic nerve in preclinical AD could be anterograde, from RGC pathology in the inner retina (where the vascular tree is sparse with low oxygen tensions (Casson et al., 2021) and amyloid deposits have been found (Koronyo et al., 2012)), or retrograde, from loss of destination cells in the thalamus and other brain targets of the optic nerve. Our study did not detect a closer association between thalamus volume and retinal measures.

We found that the pattern of subcortical associations was widely distributed. This is notable because it could be hypothesised that retinal neuronal degeneration would be primarily reflective of neurodegeneration in the areas which are most closely connected to the retinal ganglion cells, i.e., the lateral geniculate nucleus or the occipital cortex. Alternatively, it could be hypothesised that regions known to atrophy in AD may be preferentially targeted by the same neuropathology as that affecting the retina, so associations could be expected between such areas, e.g., the hippocampus, and retinal thickness. Such specific hypotheses would, however, result in missing any broader patterns of associations. Although these specific areas were associated with lower retinal thickness, our findings suggest that links between the retina and the brain in midlife are not restricted to particular regions, but that retinal neuronal measurements from OCT are instead reflective of overall brain integrity. In addition, we found evidence of targeted associations in the hippocampus, cerebellum, and ventral diencephalon that were independent of globally smaller total brain volume. Future studies should thus consider whether global or regional effects are driving any associations with retinal thickness, including whether global or region-specific effects discriminate between neurodegenerative disorders.

Limitations of this study are that the analyses conducted were cross-sectional, as MRI and OCT were conducted during the age 45 assessment. It is likely longitudinal studies with repeated OCT and MRI measures are required to determine the nature of this association. Furthermore, the sample is predominantly New Zealand European, so whether these findings generalise to people of non-white ethnicities is unknown. We were unable to confirm a diagnosis of preclinical AD in this cohort through established biomarkers, such as amyloid beta (Aβ_40_, Aβ_42_) or hyperphosphorylated tau. There are other explanations for associations between retinal thinning and structural brain integrity which do not presuppose that a person will develop AD, such as other neurodegeneative diseases, substance or alcohol abuse, or traumatic brain injury. As this is a population-based cohort, we did not exclude any participants on the basis of health status, except for those with diseases directly affecting the retinal layers or MRI brain measurements.

The overarching goal of this paper was to provide further evidence for the potential of OCT as a clinically-useful screening tool for identifying preclinical disease in the community, particularly those at risk of developing AD, and for monitoring disease progression. These findings should inform applications to improve screening of AD risk at an early stage of the disease and to ensure equitable access to such screening through the use of existing retinal imaging technology. OCT technology is already widely available in eye clinics, some primary care facilities, and most retail optometrists. Work is progressing on using OCT images in artificial intelligence/machine learning applications for the diagnosis of AD, and manufacturers of commercial ophthalmology imaging tools are likely to implement models into their devices when evidence of efficacy is clear. Potential incorporation of machine learning into OCT technology would widen the availability of retinal AD screening to regional and marginalised populations, as well as establishing and expanding large and validated normalised databases to which each individual can be compared.

In our population-based sample of middle-aged adults, we found that thinner retinal neuronal layers (both mean RNFL and mean GC-IPL) were associated with alterations in structural brain integrity associated with increased risk for later AD. These associations were evident at the whole-brain, regional, and parcel levels, suggesting that RNFL and GC-IPL reflect widespread patterns of at-risk brain structure, rather than preferentially in regions involved in the visual pathway. These findings suggest that RNFL and GC-IPL may be readily-measured indices of brain health, further supporting their possible adoption as potential early biomarkers of later AD risk.

## Acknowledgements

We thank the Dunedin Study members and their families and friends for their long-term involvement. We also thank the members of the Advisory Board for the Dunedin Neuroimaging Study, all Unit research staff, and Dunedin Study founder Dr Phil A. Silva. The Dunedin Multidisciplinary Health and Development Research Unit is based at the University of Otago within the Ngāi Tahu tribal area whom we acknowledge as first peoples, tangata whenua (trans. people of this land).

